# Strain-level differences in *Gardnerella* urinary tract persistence and pathogenesis are consistent with comparative phylogenomic analyses

**DOI:** 10.1101/2025.09.19.677092

**Authors:** Lokesh Kumar, Sonia N. Whang, Robert F. Potter, Nicole M. Gilbert

## Abstract

**Background:** *Gardnerella* is a genus of gram variable anaerobic bacteria that is commonly present in the female urogenital tract, especially during bacterial vaginosis (BV). BV is linked with increased risk of urinary tract infections (UTI) and *Gardnerella* has been frequently detected in urine collected directly from the bladder. Understanding the contribution of *Gardnerella* to urogenital pathogenesis has been complicated by its genetic heterogeneity and a shortage of data from *in vivo* models. Recently, a clinical isolate of *Gardnerella* displayed covert pathogenesis in a mouse urinary tract inoculation model, triggering urothelial exfoliation and promoting UTI by uropathogenic *E. coli*. Data from clinical studies suggests differential association of *Gardnerella* phylogenetic clades with BV or urogenital infections. *In vitro* data has demonstrated heterogeneity in the presence and expression of putative virulence determinants between *Gardnerella* strains. This study was designed to compare diverse *Gardnerella* strains *in vivo* to identify genomic variation associated with urinary tract persistence and pathogenesis.

**Methods:** Eighteen *Gardnerella* clinical isolates from each of the four main phylogenetic clades were individually inoculated transurethrally into female C57BL/6 mice. Bacteriuria was monitored by quantitative culturing of *Gardnerella* in urine. Pathologic features were assessed by immunofluorescent and histological staining of bladder tissues. Pan-genome phylogenetic analyses were performed on the 18 *Gardnerella* isolates used for mouse infections to identify accessory genes that were associated with observable *in vivo* phenotypes, including long and short-term persistence, urothelial exfoliation and bladder edema. Genes that were significantly associated to phenotype were then matched against a pangenome analysis of 291 publicly available *Gardnerella* genomes to determine the conservation of these putative colonization and virulence factors across the genus.

**Results:** *Gardnerella* strains displayed clear differences in persistence and pathogenesis in the mouse bladder that were congruent with phylogeny. Clade 2 strains were more persistent in the urinary tract whereas strains from the other three clades either caused transient bacteriuria or were undetectable. Strains from clade 2 and 4 induced urothelial exfoliation while edema was triggered by strains from clades 2, 3 and 4. Pangenome analyses revealed 45 genes that were associated with *in vivo* persistence and pathogenicity. Among the wider 291 publicly available genomes, clade 2 strains encoded more of the genes associated with bacteriuria phenotypes compared to strains in the other three clades. Exfoliation-associated genes were present in most clade 4 strains. Clade 3 strains lack most of the *in vivo*-associated genes, whereas clade 1 strains were more heterogenous.

**Conclusions:** This study provides *in vivo* evidence for differential urinary tract colonization and pathogenesis by strains from different clades/species within the genus *Gardnerella* and identifies new putative persistence and virulence factors. Utilizing the *in vivo* data from tested strains, pangenome analyses predicts that clade 2 *Gardnerella* are the most likely to persist in the urinary tract and that clades 2 and 4 have the highest uropathogenic potential. These findings inform future targeted screening and treatment approaches aimed at limiting harmful *Gardnerella* urinary tract exposures

## Introduction

*Gardnerella* is a facultative gram variable anaerobe with a gram-positive cell wall that resides within the human urogenital microbiome. The history of *Gardnerella* taxonomy and nomenclature reflects the evolving understanding of the bacterium, its role in human health, and advancements in microbiological classification. *Gardnerella vaginalis* was first described in 1955 by Gardner and Dukes, who linked it to bacterial vaginosis (then “nonspecific vaginitis”) and named it *Haemophilus vaginalis* due to its vaginal origin, heme requirement, and Gram-variable staining. Later taxonomic revisions, supported by biochemical, genetic, and ultrastructural evidence, led Greenwood and Pickett (1980) to establish the new genus *Gardnerella*, with *G. vaginalis* as the sole species. Molecular analyses, including DNA hybridization and 16S rRNA sequencing, subsequently confirmed its placement in the family *Bifidobacteriaceae* (phylum *Actinobacteria*), distinguishing it from *Haemophilus* and *Corynebacterium*. Early molecular typing divided *Gardnerella* into distinct 8 and then 17 biotypes ^1,2^. Whole-genome sequencing and comparative genomics revealed significant genetic diversity among *Gardnerella vaginalis* strains, leading to their division into four distinct clades ^3,4^. Genomic analyses, including average nucleotide identity (ANI) and phylogenomic studies, have identified at least 13 distinct genomic species within what was previously called *Gardnerella vaginalis*. In 2019, Vanechoutte et al. proposed splitting *Gardnerella* into multiple species, including *G. vaginalis, G. piotii, G. leopoldii*, and *G. swidsinskii*, based on genetic divergence and ecological differences. Since then, *G. pickettii* and *G. greenwoodii* have also been added ^5^.

Much like the history of its nomenclature, our understanding of the relationship between *Gardnerella*, health, and disease has evolved since its initial assignment as the causative agent of BV. The presence of *Gardnerella* in the vagina, even at high levels, is not always associated with symptoms. This observation has led to a prevailing hypothesis that different clades or strains of *Gardnerella* are more or less pathogenic than others. This idea is supported by *in vitro* data. Phenotypic variation has long been observed among *Gardnerella* isolates in biochemical activities (hence, early the division into “biotypes”) and expression of putative virulence features (e.g., hemolysis, biofilm formation, sialidase activity, epithelium cytotoxicity and immune activation). Studies have reported clade-wise patterns, with certain clades exhibiting robust biofilm-forming capacity, elevated sialidase activity, and strong adhesion to epithelial cells—traits that may enhance epithelial colonization and contribute to recurrent infections. In contrast, other clades demonstrate comparatively lower pathogenic potential ^6-8^. Clade-specific patterns in antibiotic susceptibility have also been reported. A recent large study found that most strains from clade 1/*G. vaginalis* and clade 2/*G. piotii/G. pickettii* exhibited sensitivity or intermediate resistance in contrast to clades 3/*G. greenwoodii* and clade 4/*G. swidsinskii/G. leopoldii* that demonstrated uniformly high-level resistance ^9^. Some clinical studies have indicated that clades 1, 2 and sometimes 3 are more associated with BV in women ^10,11^ than clade 4, while a different study found clade 4 more associated with BV ^12^. Taken together, these *in vitro* and clinical studies indicate that pathogenic heterogeneity exists among *Gardnerella* strains, but this conclusion has not been demonstrated in *in vivo* models that can establish causation.

Most research on *Gardnerella* has focused on its role in BV in the vagina and its associations with reproductive tract health. However, mounting evidence indicates that *Gardnerella* can also reach the bladder. Reports of the isolation of *Gardnerella* from urine collected directly from the bladder by suprapubic aspiration go back more than 50 years. McFadyen and Eykyn (1968), McDowall *et al*. (1981), and Savige (1983) reported the isolation of *Gardnerella* from 15.9%, 18% and 22%, respectively, of suprapubic aspirated (SPA) urine from >1,000 pregnant women ^13-15^. Because bacteriuria had disappeared in many of their patients when SPA was repeated later in pregnancy and pyuria was rarely detected, McFadyen and Eykyn believed that the presence of *Gardnerella* in urine was not significant. However, they reported an increased incidence of symptomatic urinary tract infection in women with detectable *Gardnerella* (13.5%) compared with non-bacteriuric women (4%). *Gardnerella* continued to be one of the most frequently cultured fastidious organism, present in ∼25% of SPA and catheterized urine samples from both pregnant and non-pregnant women in multiple studies over the next decade ^15-18^. The expanding urobiome field continues to routinely detect *Gardnerella* by culture and molecular techniques in catheterized and clean-catch urine samples from women ^19-21^. The frequent detection of *Gardnerella* in SPA and catheterized urine samples suggests its ability to colonize the bladder, either transiently or persistently ^22^. *Gardnerella* is rarely the cause of UTI ^23^. However, results from mouse models have demonstrated that a clade 2 strain (8151B) of *Gardnerella* behaves as a “covert pathogen” capable of promoting UTI by uropathogenic *E. coli* (UPEC). *Gardnerella* 8151B was cleared from the mouse urinary tract within 12 h, but nonetheless initiated urothelial exfoliation and induced a transcriptional signature of urothelial turnover and inflammation ^24,25^. Subsequent inoculation of UPEC into *Gardnerella*-exposed bladders resulted in more persistent UPEC infection compared to mice not exposed to *Gardnerella* ^25^. In another model, *Gardnerella* 8151B inoculation into mice harboring quiescent intracellular reservoirs of UPEC from a previous infection triggered re-emergence of UPEC to cause recurrent UTI ^24,26^. These data provide one explanation for the association between BV and UTI observed in women ^27-30^. Beyond a potential connection to UTI, the presence of *Gardnerella* in urine has been associated with other urologic conditions, including urgency urinary incontinence (UUI) ^20,21^.

The direct impact of urogenital bacteria such as *Gardnerella* on the bladder mucosa and UTI outcomes, including the effects of transient exposures and strain-level phenotypic heterogenicity has remained largely unexplored. In this study, we utilized an established mouse model of urinary tract inoculation to discriminate between colonizing, exfoliating, and inflammatory *Gardnerella* strains in the bladder. We then applied comparative genomics methodologies to determine the association of pathogenic features *in vivo* with accessory gene presence. The link between bacterial genes and pathogenic traits can reveal new hypotheses regarding *Gardnerella* tropism and virulence for future mechanistic studies. This is particularly important for *Gardnerella* as a tractable reverse genetics system to study genetic contributions to virulence does not currently exist. Additionally, if *Gardnerella* analysis identifies a genetic signature of uropathogenesis, this could serve as a clinical biomarker for guiding future patient screening and treatment approaches. This systematic effort to delineate the clade and strain-specific phenotypes of *Gardnerella in vivo* opens the door for future studies to delineate mechanisms of *Gardnerella* urogenital pathogenesis.

## Material and methods

### Mice strains

Female C57BL/6 mice, aged 6-7 weeks old were procured from Jackson Laboratories (000664) and housed in a specific pathogen-free (SPF) facility at Washington University School of Medicine and maintained under controlled environmental conditions, including a 12-hour light/12-hour dark cycle, with unrestricted access to standard chow and water. All of the mice were allowed for acclimatization at least one week before starting the experiments. Mouse experiments were carried out in strict accordance with the recommendations in the Guide for the Care and Use of

Laboratory Animals. The Institutional Animal Care and Use Committee (IACUC) of Washington University School of Medicine, St. Louis, MO, USA, approved all procedures in advance (Protocol Numbers: 23-0015).

### *Gardnerella* strains and growth conditions

A total of eighteen *Gardnerella* strains from clades 1, 2, 3, and 4 were used in this study as listed in Table 1. Strains were streaked from glycerol freezer stocks onto NYCIII agar plates (Lewis et al 2013) and grown at 37 °C under in an anaerobic chamber (Coy Laboratory, Grass Lake, MI, USA) for 24-48 hrs. Strains were then subcultured from these plates in NYCIII Broth grown for 24 h at 37 °C under anaerobic conditions. Bacterial inocula were prepared in PBS in the anaerobic chamber.

**Table 1.**
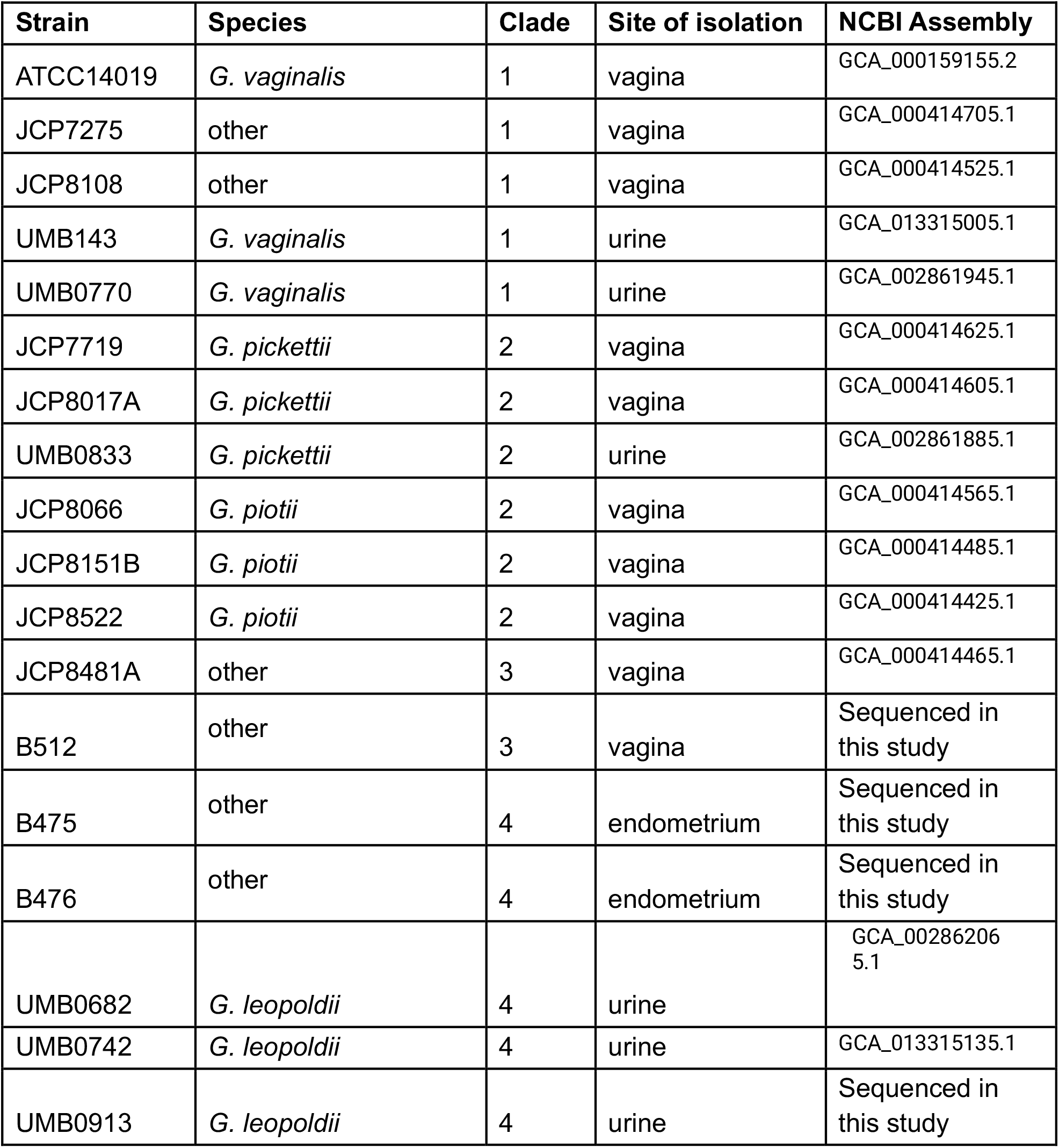
*Gardnerella* strains used in this study.

### *Gardnerella* infection in mouse model

To evaluate the colonization dynamics in the urinary tract, *Gardnerella* inoculations were performed in mice (**Fig 1A**) Mice were anesthetized using isoflurane and 50 µL *Gardnerella* inocula containing 1 × 10^8^ colony forming units (cfu) were transurethrally inoculated, following established protocol ^31^. Each experimental and mock group consisted of at least five mice. Eighteen hours post-inoculation, urine samples were collected from each mouse to quantify bacterial titers by serial dilution and plating on NYCIII in the anaerobic chamber. Subsequently, mice received a second transurethral inoculation with either the same *Gardnerella* strain or phosphate-buffered saline (PBS) for mock controls. Six hours after the second exposure (24 hours after the initial inoculation), urine was collected for cfu enumeration, and all mice were humanely euthanized via cervical dislocation under isoflurane anesthesia. Bladders were aseptically harvested for further studies. As a positive control for urothelial exfoliation, a separate group of mice was inoculated with chitosan as described previously ^32^.

**Figure 1.**
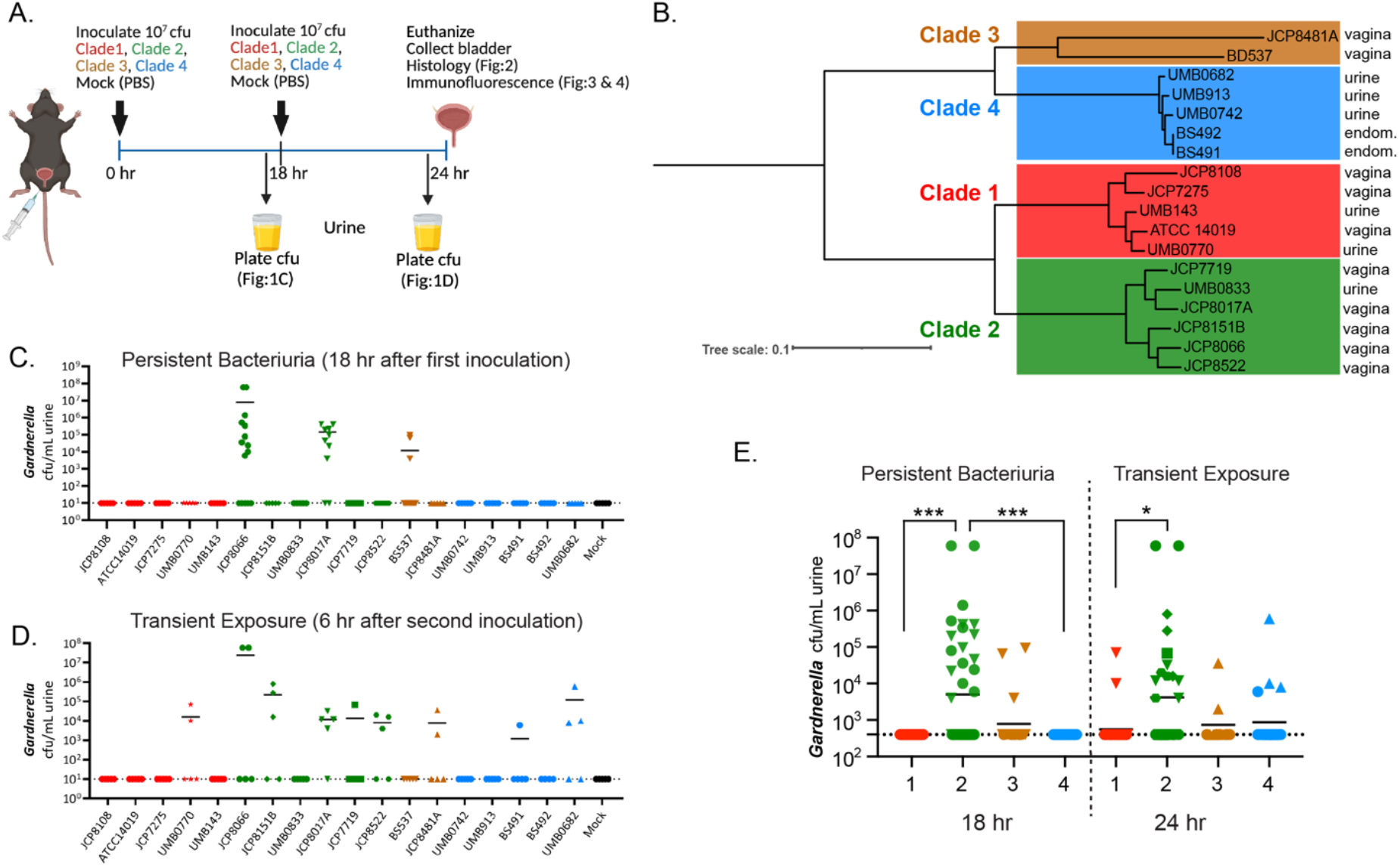
Differences in urinary tract clearance rates between *Gardnerella* strains: (A) Schematic representation of the experimental timeline illustrating *Gardnerella* exposure in the urinary bladder of a C57BL/6 mouse model. The diagram includes time points for transurethral inoculation and subsequent urine collection for cfu. (B) Phylogenetic tree depicting the 18 *Gardnerella* strains tested *in vivo* in this study. The four main clades are represented by a unique color: clade 1-red, clade 2-green, clade 3-brown, and clade 4-blue. Site of isolation of each strain is listed on the right. (C) *Gardnerella* cfu in urine samples collected 18 hours after the first exposure. Each dot represents an individual mouse. (n=5 per strain, except for strains JCP8066 (n=15) and JCP8017A (n=10). (D) *Gardnerella* cfu in urine samples collected 24 hours after two sequential exposures (6 hrs. after second inoculation). Each dot represents an individual mouse (n=5 per strain). (E) Bacteriuria data from panels C and D, combined according to *Gardnerella* clade.

### Histological and Immunofluorescence Analysis of Mouse Urinary Bladder Tissues

Mouse urinary bladders were collected 24 h post infection and fixed overnight in 4% paraformaldehyde and then transferred to 70% ethanol. The bladders were embedded in paraffin, and sagittal sections were prepared then mounted on glass slides by Nationwide Histology (Missoula, MT). Slides were deparaffinization using xylene and then moved to isopropyl alcohol for 5 mins three times and then placed under running water for 10 min. For histological analysis, slides were stained with hematoxylin and eosin (H&E) according to standard protocol. Sections were observed on an Olympus BX40 microscope equipped with a mounted Zeiss Axiocam 305 camera. The presence/absence of edema in each bladder section (4-5 bladders per experimental group) was determined by two independent observers who were blinded to the experimental group.

For immunofluorescence analysis, deparaffinized and rehydrated sections underwent antigen retrieval by boiling in pH 9 antigen retrieval buffer (Vector Laboratories, H-3301) for 30 minutes. After cooling to room temperature, sections were permeabilized with 0.5% Triton X-100 in phosphate-buffered saline (PBS) for 15 minutes. Non-specific binding was blocked by incubating sections in blocking buffer containing 5% bovine serum albumin (BSA) in 0.5% Triton X-100 PBS for 1 hour at room temperature. Primary antibodies Cytokeratin-20 (CK20; Proteintech,17329-1-AP) and e-cadherin (R&D system, AF748) were applied to the sections, and slides were incubated overnight at 4°C in a humidified chamber. The following day, sections were washed three times with 0.5% Triton X-100 PBS and incubated with appropriate fluorophore-conjugated secondary antibodies for 1 hour at room temperature. After washing, nuclei were counterstained with DAPI, and slides were mounted using an antifade reagent. Fluorescent images were acquired using a Zeiss Axio Imager M2 Plus microscope. The presence/absence of exfoliation as evidenced by loss of CK20 staining of the lumenal surface of the urothelium in each bladder section (4-5 bladders per experimental group) was determined by two independent observers who were blinded to the experimental group.

### Whole-genome sequencing and assembly

Glycerol stocks of the four *Gardnerella* isolates sequenced de novo in this study (B512, B476, B475, and UMB0913) were subcultured onto NYCIII agar plates and grown under anaerobic conditions at 37 °C for 24-48 h. A loopful of colonies was collected and washed in sterile PBS and then resuspended in DNA/RNA Shield (Zymo Research Corporation, Tustin, CA). The samples were sent overnight to Plasmidsaurus (South San Francisco, CA, USA) for bacterial Whole-genome sequencing with extraction. Accordingly, as described on their website (https://plasmidsaurus.com/products/bacterial-genome), genomic DNA is processed for Oxford Nanopore Technology sequencing using v14 library prep chemistry and R10.4.1 flow cells. Raw Oxford Nanopore reads were quality-filtered by removing the lowest 5% using Filtlong v0.2.1 ^33^. Reads were down sampled to 250 Mb for an initial draft assembly with Miniasm v0.3 ^34^. Based on the Miniasm output, reads were re-downsampled to ∼100× coverage (or retained in full if coverage was lower) with increased stringency against low-quality reads to preserve small plasmids. The final assembly was generated with Flye v2.9.1, followed by polishing with Medaka v1.8.0 ^35^. Assembly from Plasmidsaurus was used for downstream analysis.

### Phylogenomic analysis

*Gardnerella* spp. genomes from NCBI Genome dataset were downloaded on May 8, 2025. Genomes derived from metagenomic DNA were excluded from the study. The resulting genomes (n=287) were included with the four de-novo sequenced *Gardnerella* genomes into a total cohort (n=291). All of the genomes had protein coding sequences annotated with bakta v1.11.3^36^.

bakta --db {database path} --output {output path} {input path} –force --locus-tag {name} -- locus{name})

The resulting GFF3 files were constructed into a core genome alignment using panaroo v1.5.2 ^37^.

panaroo --i {GFF path} --o {output directory} -a core -c.8 -f.5 --len_dif_percent.9 --remove-invalid-genes -t 12 --clean-mode strict

The alignment was converted into an approximate maximum likelihood phylogenetic tree with FastTree and viewed in the iTOL v6 webserver^38,39^.

fasttree -nt -gtr -gamma {path to core_gene_alignment_filtered.aln} > {output path}

This process was repeated on a separate folder containing just the GFF3 files from the 18 experimentally used strains. Next, we used scoary 1.6.16 to identify accessory genes (default command) that are associated with the presence of specific experimentally investigated strains (exposure, exfoliation, edema, and colonization) in the 18 cohort panaroo analysis ^40^. Only outputs showing a positive association from scoary (e.g. genes with sensitivity >.5 were included in analysis). Next, we matched the genes with the significantly associated to phenotype against all isolate panaroo analysis. The pan_genome_reference.fa file produced by panaroo was uploaded to the eggNOG-mapper v2 webserver ^41^. Four of the genes were excluded by panaroo in the all-genome analysis, however 41/45 of the phenotype associated genes were linked to their counterpart in the panaroo-all dataset. These genes were portrayed as a presence/absence matrix against the Clade classified phylogenetic tree in iTOL.

### Data availability

The raw fastq files for the four *Gardnerella* genomes sequenced *de-novo* in this study are uploaded to NCBI BioProject PRJNA1304581.

## Results

### Differences in urinary tract clearance rates between *Gardnerella* strains

We previously reported that a clade2/*G. piotii* strain JCP8151B did not colonize the urinary tract of female C57BL/6 mice beyond 12 h. Urothelial disruption and augmentation of UPEC UTI required two transurethral inoculations ^24,26,31^. Here we used the same two-exposure model and experimental timeline to examine *Gardnerella* colonization patterns and pathogenesis in the urinary tract (**Fig 1A**). We used 18 *Gardnerella* clinical isolates collected primarily from vaginal swabs or urine and two endometrial isolates. These strains were distributed across the four main clades, albeit with the least representation from clade 3, which is also the rarest among published *Gardnerella* genomes and isolates (**Fig 1B**). Mice were transurethrally inoculated with ∼10^8^ colony forming units (cfu) of a single *Gardnerella* strain. Urine was collected 18 hours post inoculation (hpi), immediately before administering a second *Gardnerella* inoculation. The urine was collected again six hours after the second inoculation (24 h after initial exposure) and then bladders were collected and fixed for histological and immunofluorescence microscopy. This timepoint was chosen because it coincides with when JCP8151B caused epithelial exfoliation and augmentation of UPEC UTI in our prior studies. Thus, 8151B served as a positive control for these experiments.

Our findings revealed distinct differences in the rates of bacteriuria between *Gardnerella* strains. Bacteriuria results could be categorized into three main phenotypes: “Persistent Bacteriuria” where cfu were detected 18 h following the first inoculation (**Fig 1C**), “Transient exposure” where cfu were detected 6 h after the second inoculation (**Fig 1D**), or no cfu detected at either timpoint. We hypothesized that strains isolated from urine could represent those better able to colonize the urinary tract. However, of the five strains that displayed high rates of bacteriuria at either timepoint, four were vaginal isolates and only one was isolated from urine. There were apparent differences in colonization at the clade level. All strains from clade 1/*G. vaginalis* and clade 4/*G. leopoldii* were cleared after the first exposure and undetectable at 18 h (**Fig 1C**). This is consistent with our previous results with JCB8151B, recapitulated here, and was true for UMB0833, JCP7719, and JCP8522 clade 2/*G. piotii* strains (**Fig 1C**). However, experiments revealed that two clade 2/*G. piotii* strains, JCP8066 and JCP8017A, displayed persistent bacteriuria following a single inoculation (11/15 and 8/10 mice, respectively; **Fig 1C**). A low rate of bacteriuia was also found for clade 3 B512 (3/10 mice; **Fig 1C**). More *Gardnerella* strains displayed bacteriuria 6 h following the second exposure (**Fig 1D**). Strains displaying a bacteriuria rate >50% included three clade 2/*G. piotii/G. pickettii* (JCB8151B, JCP8017A, JCP8522) and one clade 4/*G. leopoldii* (UMB0682) (**Fig 1D**). Low but detectable bacteriuria rates were observed for one Clade 1/*G. vaginalis* (UMB0770) one Clade 3 (JCP8481A) one Clade 4 (B475) strains (**Fig 1D**). Of the strains that displayed bacteriuria at 18h, JCP8017A displayed a high rate of bacteriuria at both time points, whereas JCP8066 and B512 were both more often cleared after the second exposure, despite the shorter length of time between inoculation and urine collection. Follow-up experiments with only a single inoculation of these three strains showed the same pattern of detectable bacteriuria at 18 h but clearance within 24 h. When examined collectively, clade 2 *Gardnerella* caused significantly higher levels of persistent bacteriuria than clade 1 and clade 4 and significantly higher transient exposure cfu than clade 1 (**Fig 1E**). The difference between clade 2 and clade 3 was not statistically significant, likely because there were fewer clade 3 strains tested. Overall, these results demonstrate strain specific differences in *Gardnerella* urinary tract colonization dynamics, with clade 2 strains collectively showing significantly higher levels of bacteriuria compared to the other clades.

### *Gardnerella* strain differences in urothelial exfoliation

The urothelium is a dynamic and multifunctional tissue essential for urinary bladder homeostasis and acts as a barrier for infection. Exposure to uropathogens leads to significant exfoliation of the bladder urothelium, that serves as a host defense protective mechanism to eliminate infected cells. Our previous studies demonstrated that two exposures to *Gardnerella* JCP8151B triggered urothelial apoptosis and exfoliation. Next, we examined whether exfoliation was triggered by more genetically diverse *Gardnerella* strains by staining Cytokeratin 20 (CK20), a marker of superficial umbrella cells in bladder sections collected in from the experiments outlined in Fig 1. As a positive control, we also included a group of mice treated with chitosan, which is known to induce urothelial exfoliation ^32,42^. Bladders from control mice that were mock infected with PBS displayed even and continuous CK20 expression localized to the cytoplasm of superficial umbrella cells lining the bladder lumen (**Fig. 2A**). Chitosan-treated mice displayed the expected loss of CK20 staining on the lumenal surface (**Fig 2A**). Urothelial exfoliation characterized by a loss of superficial CK20 staining was also evident in the bladders of some mice inoculated with *Gardnerella* (**Fig 2A and Fig S1, denoted with +**). The proportion of mice in each group that displayed exfoliation (exfoliation rate) was determined for each strain and compared to mock infected controls by Fisher’s exact tests (**Fig 2B**). *Gardnerella* strains could be divided into two categories. ‘Non-exfoliators’ displayed healthy urothelial staining pattern with no significant difference from mock controls, whereas ‘Exfoliators’ had significantly higher exfoliation rates. There was no obvious relationship with isolation site as vaginal and urine isolates were present in both categories. Most clade 1/*G. vaginalis*, except for JCP8108, and all clade 3 strains were non-exfoliators. Most clade 2/*G. piotii/G. pickettii* strains, except JCP8017A and UMB033, and all clade 4/*G. leopoldii* strains were exfoliators. When analyzed collectively, clade 2 and clade 4 *Gardnerella* induced a significantly higher exfoliation rate compared to controls (**Fig 3C**). These data demonstrate that there is strain variability in urothelial damage induced by *Gardnerella*, with clades 2 and 4 showing the highest exfoliation rates.

**Figure 2.**
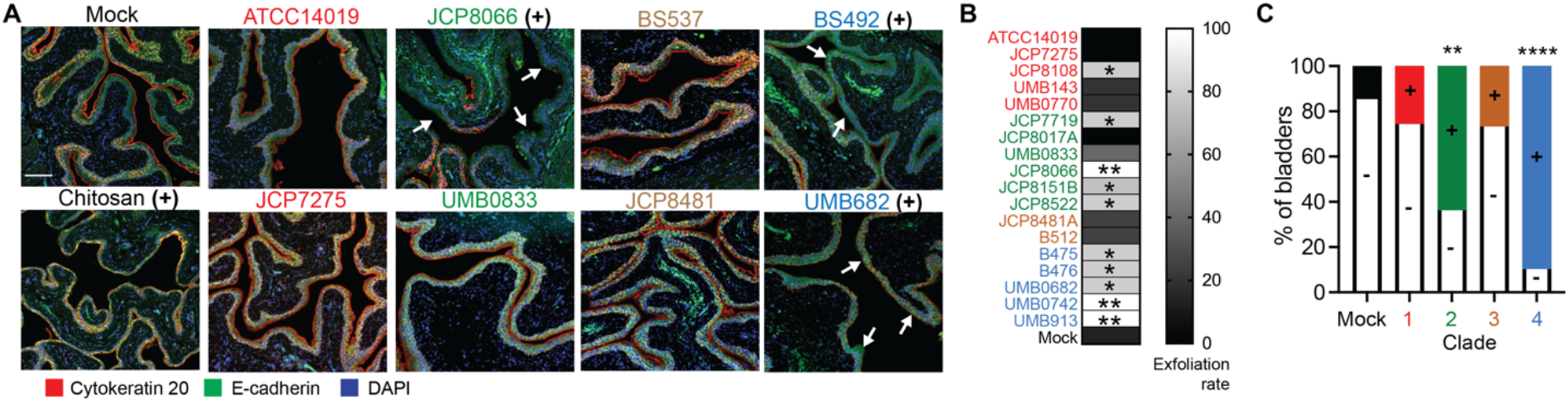
Strain differences in urothelial exfoliation induced by *Gardnerella*. (A) Bladders collected from mice inoculated with the indicated strains, PBS (Mock) or chitosan and stained for cytokeratin-20 (AF647, red; marker of terminally differentiated superficial umbrella cells), E-cadherin (AF488, green) and nuclei (DAPI, blue). Regions of exfoliation are denoted by white arrows. Images are shown for two strains representative of the phenotype from each clade. Scale bar = 100 μm. (+) denotes “Exfoliator” strains based on the statistical analysis shown in panel B (B) Heat map representation of the exfoliation rate calculated based on the proportion of mice in each group that displayed exfoliation. C) Exfoliation data from tested *Gardnerella* strains combined according to clade. Colored (+) bars denote the percentage of bladders with exfoliation. * P < 0.05, ** P < 0.01, **** P < 0.0001, Fisher’s exact tests.

**Figure 3.**
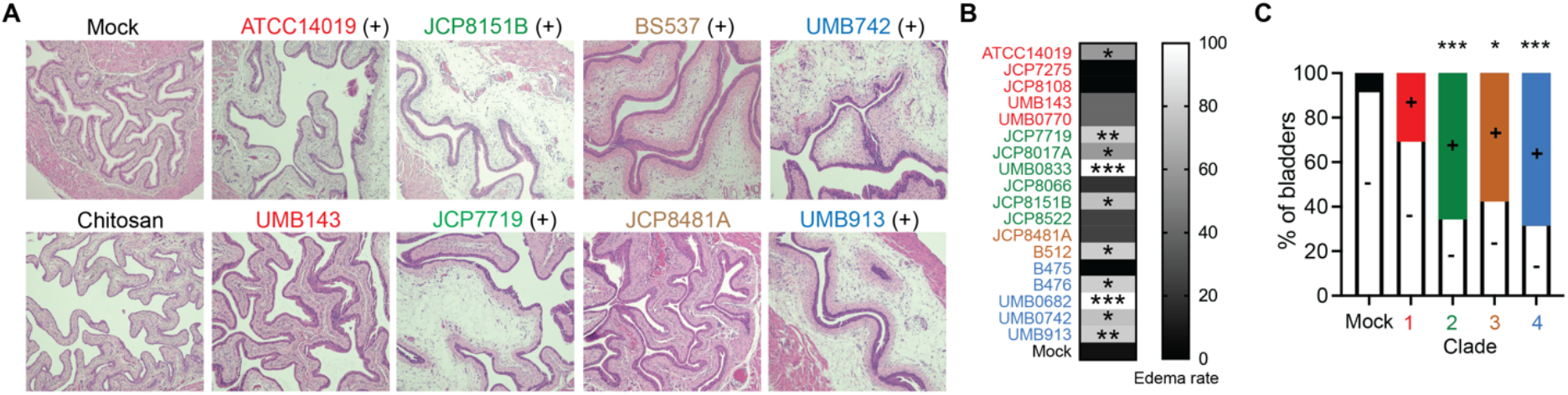
Strain differences in *Gardnerella* induction of bladder edema: (A) Representative hematoxylin and eosin (H&E)–stained sections of mouse urinary bladder tissues collected after two transurethral exposures of *Gardnerella*. Panels display bladder sections from Mock (PBS), Chitosan-treated mice (a transient urothelial disruptor), and for two strains representative of the phenotype from each clade. Each clade is represented by a different color scheme: clade 1 (red), clade 2 (green), clade 3 (brown), and clade 4 (blue). Scale bar = 100 μm. (+) denotes “Edema” strains based on the statistical analysis shown in panel B (B) Heat map representation of the edema rate calculated based on the proportion of mice in each group that displayed bladder edema. (C) Edema data from tested *Gardnerella* strains combined according to clade. Colored (+) bars denote the percentage of bladders with edema. * P < 0.05, ** P < 0.01, *** P < 0.001, Fisher’s exact tests.

### Some *Gardnerella* strains cause edema and inflammation in the bladder

During our analysis of urothelial exfoliation, we noticed that some bladders displayed edema in the lamina propria (**Fig 3A and Figure S2**). Edema is a key component of the acute inflammatory host response to uropathogen infection in the urinary bladder. It is characterized by fluid accumulation in the interstitial spaces accompanied by infiltration of leukocytes. *Gardnerella* could be divided into two categories based on our blinded analysis of bladder histology. ‘No edema’ strains displayed healthy urothelium without histological inflammation in the majority of exposed bladders. ‘Edema’ strains induced notable edema in the majority of exposed bladders. To our surprise, more than half of the tested *Gardnerella* strains induced edema (**Fig 3B**). Most of the clade 1 strains, with the exception of ATCC14019, did not cause edema. Cumulatively, clades 2, 3 and 4 displayed a statistically significant increased rate of edema compared to mock (**Fig 3C**). Four of six clade 2/*G. piotii/G. pickettii* strains, one of two clade 3 strains, and 4 of 5 clade 4/*G. leopoldii* strains caused edema. When comparing the edema and exfoliation results at the strain level, there was not an obvious connection between the two phenotypes. Clade 1/*G. vaginalis* ATCC14019 and clade 3 B512 caused edema but not exfoliation. Clade 1/*G. vaginalis* JCP8108 and clade 2/*G. piotii* JCP8066 and JCP8522 caused exfoliation but not edema. There was also not a clear relationship between either phenotype and persistence in urine. All strains that were detectable in urine (**Fig 1**) triggered either exfoliation or edema. However, the converse was not true. Some strains triggered either or both phenotype but were not detectable in urine. In summary, the results from our mouse inoculation studies demonstrated clear strain differences in *Gardnerella* persistence and pathogenesis in the bladder.

### Accessory genome linkage to phenotype

We hypothesized that strain level differences we observed in our mouse model could be traced to genetic differences between the isolates. We initially performed pan-genome phylogenetic analysis on the 18 *Gardnerella* isolates used for murine infections. We performed four comparisons, based on our results for persistence (**Fig 1**), exposure (**Fig 1**), exfoliation (**Fig 2**), and edema (**Fig 3**). We determined that of the 2,334 sized pan-genome containing 1,672 accessory genes, only 2.7% (45/1672) genes are positively associated with a phenotypic response (**Fig 4**) (**Table S1**). The bacteriuria phenotypes had the most genes, with 34 associated with exposure and 7 associated with persistence. One gene (group_1083, NCBI Protein: CDC16457.1) was associated with both persistence and exposure. Exfoliation and edema were each associated with two genes. Only 16/45 of the genes had a Cluster of Orthologous (COG) assigned to their function. The most prevalent COG (4/45) was M: Cell wall/membrane/envelope biogenesis. Most of the genes (30/45) have a predicted annotation available that is more specific than “hypothetical protein”. Of these, many could be important for the host-microbe interface, either due to annotations linked to the cell wall or prior data implicating them in virulence. For example, three genes are annotated as containing an LPxTG motif, positing that they are extracellular facing via sortase activity. Notably, *vly*, the gene that encodes the toxin vaginolysin was significantly associated with edema. An additional five genes have annotations consistent with a role in carbohydrate metabolism – including hydrolysis (type I pullulanase, sialidase domain containing protein), modification (GHMP kinase C-terminal domain-containing protein), binding (Extracellular matrix-binding protein ebh GA module domain-containing protein), and synthesis (Glycosyltransferase involved in cell wall biosynthesis). Additional genes implicated in the host-microbe interface include Rib/alpha-like domain containing proteins, which form an immunoglobulin fold and were important for virulence in *Streptococcus* spp ^43^. The remaining genes with annotations that are intracellular do not have an immediate connection to the host-microbe interface but could alter global responses. These include ribonuclease III, toxin-antitoxin system antitoxin component Xre domain protein, and HTH merR-type domain-containing protein.

**Figure 4.**
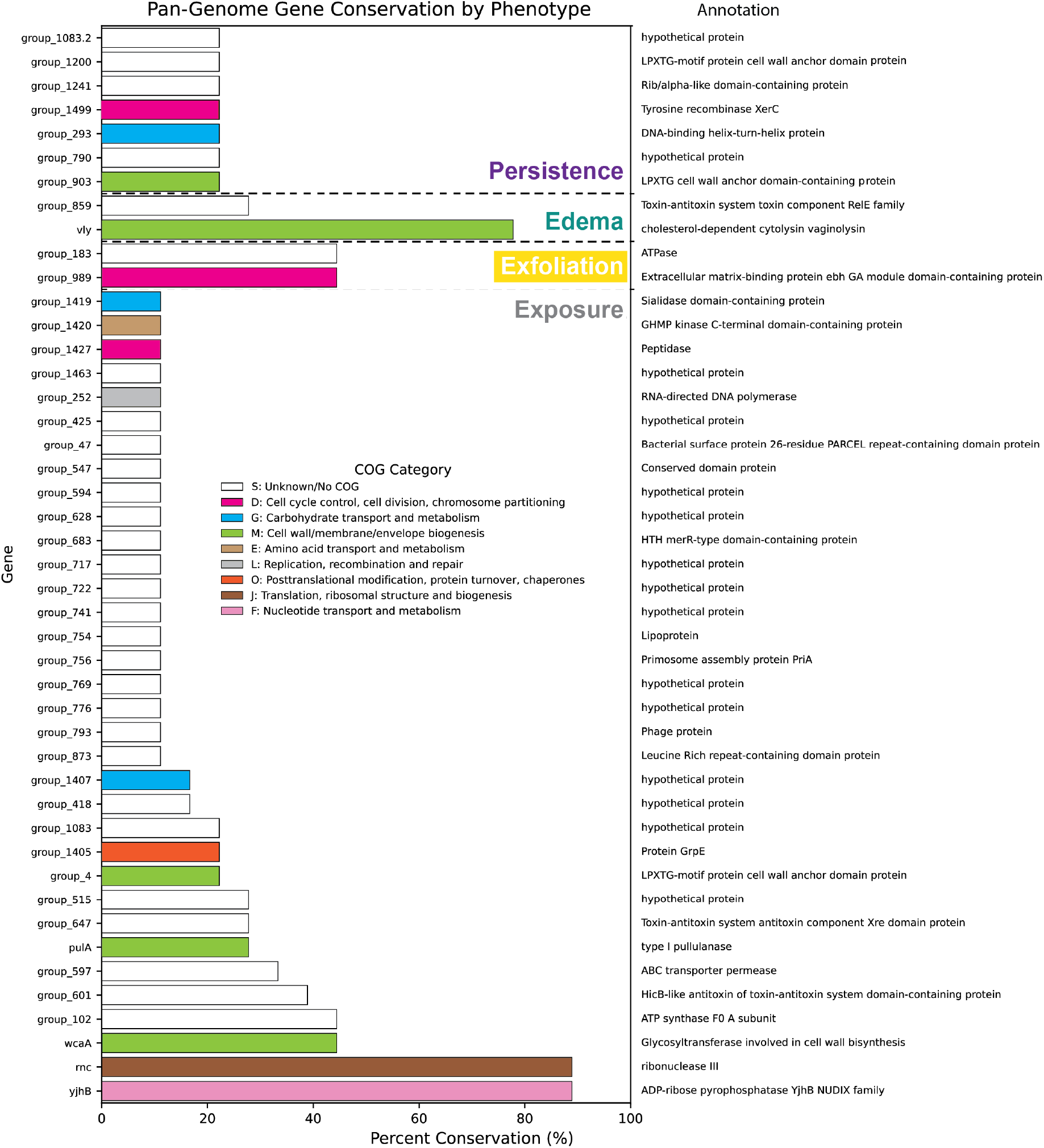
Visualization of scoary identification of accessory genes significantly associated with phenotypic traits. Panaroo gene identifications are on the far left and annotations are on the far right. Barplot depict the percent conservation within the 18 isolate cohort used for transurethral inoculation. Colors depict COG categories determined by eggNOG-mapper v2.

### Conservation of virulence-associated genes within *Gardnerella* genus

Results from mouse infection experiments revealed clade and strain-specific difference in colonization and uropathogenesis by *Gardnerella*. The results from our focused pan-genome analysis suggested that *Gardnerella* strains displaying urinary tract persistence or pathogenesis have distinct accessory gene profiles. Given that *Gardnerella* is now recognized as a genus containing multiple species within a variety of clades, we next determined the conservation of these putative virulence factors across the genus. To address this, we performed pan-genome analysis on a cohort of 291 publicly available *Gardnerella* genomes from all four clades; 39% (114/289) were clade 1, 17% (49/291) were clade 2, 10% (29/291) were clade 3, 30% (86/291) were clade 4, and 4% (13/291) were not in one of the four clades (**Table S2**). The total size of the pan-genome was 4,579 genes with 455 in the core-genome. Placement of the 18 genomes used for functional analysis found all of them within their expected clade (**Fig 5**). Our results reflect the variation in accessory genome composition that has previously been described within *Gardnerella*. Four of the 45 genes that were associated with *in vivo* phenotypes were not found in the larger pan-genome output, three from exposure and one from colonization. Group_278, annotated as containing an LPxTG motif was the most conserved colonization associated gene, found in 59% (171/291) genomes. Four genes were found in <10 isolates. Vly, the gene encoding the toxin vaginolysin associated with edema was the 3^rd^ most conserved gene overall, and was found in 90% (262/291) of all *Gardnerella* genomes with absence in a portion of Clade 1 and Clade 2 genomes. Clade 2 strains tended to encode more of the genes associated with *in vivo* persistence (purple) and exposure (gray) compared to strains in the other three clades. Consistent with the results of our tested strains, Clade 3 strains tended to lack most of the *in vivo*-associated genes. A substantial majority of clade 4 strains encoded both exfoliation-associated genes, with their presence lacking in only eight clade 4 strains. This pattern echoes the *in vivo* results and suggests that induction of exfoliation extends broadly across clade 4 *Gardnerella*.

**Figure 5.**
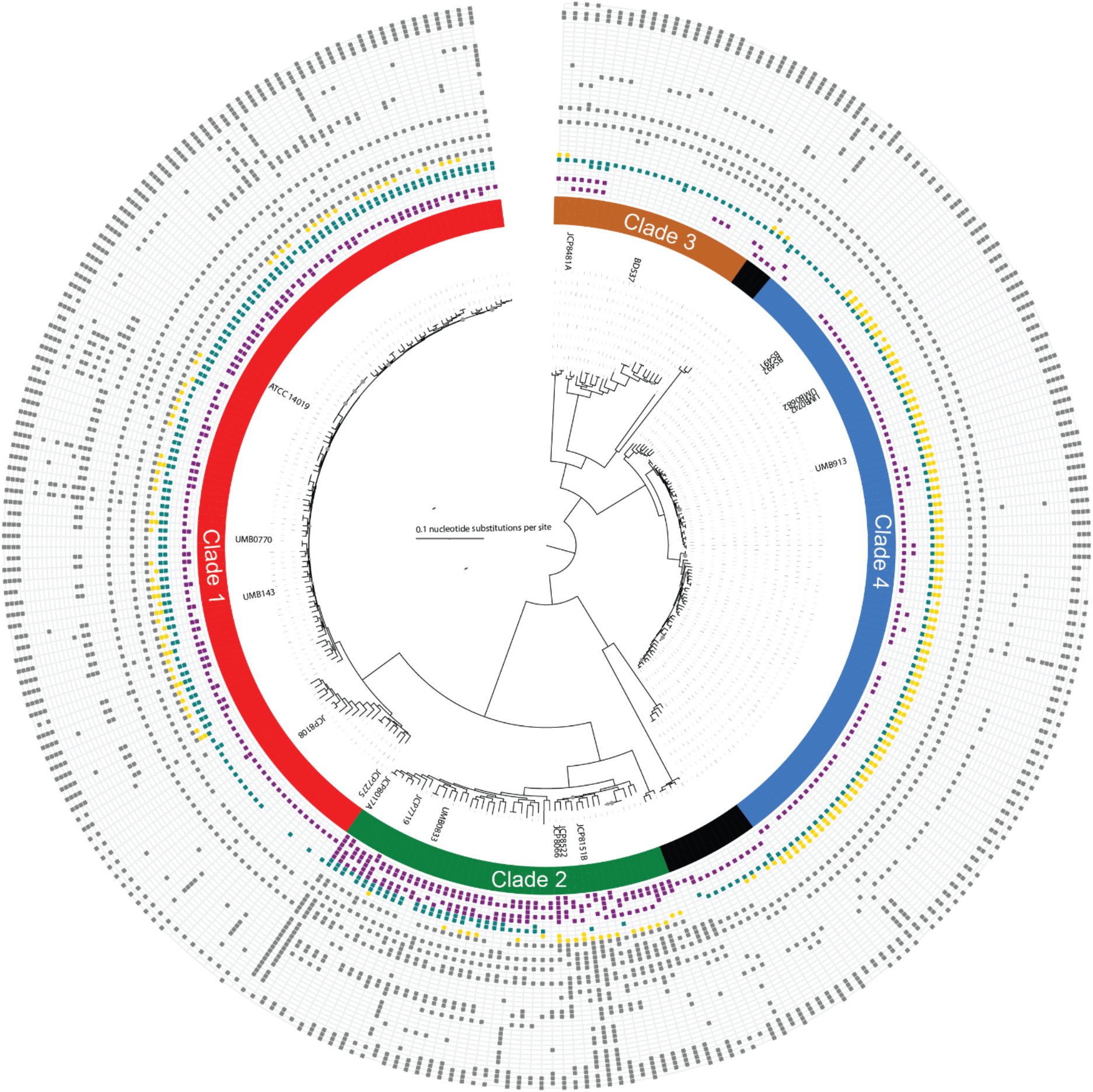
Conservation of genes associated with *in vivo* phenotypes across the *Gardnerella* genus. Core-genome phylogenetic tree depicting the relatedness of 291 publicly available *Gardnerella* genomes. The 18 strains used for transurethral infection have their strain names depicted adject to their edge of the tree. Clade identity is depicted as an inner-ring of the tree. Outer edges of the tree depict the presence (rectangle) or absence (no rectangle) of the identified scoary hits within the broader *Gardnerella* pan-genome. Purple-persistence, teal-edema, gold-exfoliation, gray-exposure. Please see **Table S2** for detailed information on gene identity corresponding to each ring.

## DISCUSSION

Urobiome studies have frequently detected *Gardnerella*, and often as the dominant organism. Thus far, urobiome studies in humans cannot distinguish whether the bacteria present in urine samples reflect stable bladder colonization or are capturing transient exposures from adjacent niches. Our previous work found that *G. piotii* strain JCP8151B was cleared rapidly from the mouse urinary tract, being undetectable in most mice at 12 hours post inoculation ^26^. Most *Gardnerella* strains tested here followed a similar pattern of rapid clearance from urine. Only two strains from clade 2, one *G. piotii* and one *G. pickettii*, showed urinary tract persistence for 18 hours following a single inoculation. Seven genes were associated with this longer-term persistence phenotype. The presence of these genes was largely clustered in clade 2 strains, but most strains only encode four or fewer. Four clade 2 strains encoded 5 persistence-associated genes and only JCP8066 and JCP84181A contained all 6 genes. Interestingly, the two strains that persisted for 18 h displayed different patterns following the second inoculation. JCP8066 was cleared following the second exposure whereas JCP8017 was detectable six hours after a second inoculation. A potential explanation for this difference could be that JCP8066 triggered exfoliation, which is believed to help eliminate uropathogens from the bladder. That said, the relationship between exfoliation and *Gardnerella* clearance was not so clear-cut since three strains that triggered exfoliation (JCP8151B, JCP8522, UMB0682) were nonetheless detectable at the six-hour timepoint. Four strains were detectable six hours after a second inoculation, and 34 genes were associated with shorter-term exposure. Again, the presence of these genes tended to cluster in the clade 2 strains, but a few of the Clade 1 strains also encoded a substantial portion. Three of the genes associated with persistence or exposure were annotated as encoding LPxTG cell wall anchor domains, indicating that they are likely surface-exposed. Futures work could explore whether these genes encode adhesins that help facilitate attachment to urothelial cells. Together our data indicate that extended urinary tract colonization may be a rare phenotype among *Gardnerella* strains.

Our previous work demonstrated that Clade 2/*G. piotii* JCP8151B triggers exfoliation of the superficial bladder urothelial cells ^25^. Here we found that this phenotype was shared by most clade 2 and all clade 4 strains tested. The shedding of urothelial cells could be crucial for the pathogenesis of UTIs, as it removes the superficial layer of the bladder epithelium that is the primary site for uropathogen adhesion and invasion. Exfoliation exposes underlying cell layers, making them more susceptible to infection. The clinical literature supports the idea that BV-associated bacteria like *Gardnerella* could promote UTI in women. Meta-analysis of four studies, analyzing a total of 1,128 women, found an increased risk of UTI in women with clinically defined BV (by Amsel criteria) when compared to without BV ^27-30^. Our previous work in mice showed that transient exposure to clade 2/*G. piotii*JCP8151B can facilitate recurrent urinary tract infections (rUTIs) by triggering *E. coli* egress from latent bladder reservoirs, a process we linked to urothelial exfoliation ^44^. Building on these insights, future studies could examine whether exfoliation could be used as a potential marker clinically to distinguish pathogenesis of *Gardnerella* in the urinary tract and associated UTI progression. Epithelial exfoliation also occurs in the vagina and increases significantly during BV ^45^. We and others have reported that *Gardnerella* triggers epithelial exfoliation and barrier disruption in mouse models of vaginal colonization ^46-48^. Direct comparison of vaginal exfoliation between different *Gardnerella* strains have not been reported, so whether strains from clades 2 and 4 are universally more exfoliating is not yet known. In addition to enhancing UTI, disruption of the urothelium and suburothelium has been implicated in the symptom of urgency associated with overactive bladder (OAB) and urgency urinary incontinence (UUI) ^49^. A few studies have linked urinary *Gardnerella* to UUI or OAB, including one that found increased *Gardnerella* in patients that displayed reduced efficacy of OAB treatment ^21,50,51^. A recent cross-sectional study found that women with BV (therefore having high vaginal *Gardnerella*) have higher rates of UUI ^52^. Future studies will examine whether *Gardnerella* affects voiding behaviors in the mouse model.

The *in vivo* studies in mice uncovered edema as a previously unknown phenotype of *Gardnerella* urinary tract exposure. Like exfoliation, edema was more frequently induced by clade 2 and clade 4 strains. Several strains caused both pathologic features, whereas strains ATCC14019 and B512 only caused edema and JCP8108, JCP8066, and B475 only caused exfoliation. These data indicate that edema did not depend on exfoliation. It could be reasoned that exfoliation and inflammation could facilitate the observed rapid clearance of *Gardnerella* from the bladder. However, we did not observe a clear relationship between the triggering of either of these phenotypes and the cfu detected in urine. The fact that the colonizer and exposure strains all triggered edema, exfoliation, or both phenotypes suggests that these strains have mechanisms for persisting in the face of these host responses and that the duration of *Gardnerella* presence in the bladder is important mediator of pathologic outcomes. The patten of edema present in *Gardnerella*-exposed bladders is histologically similar to that observed in a cyclophosphamide (CYP)-induced interstitial cystitis (IC) mouse model ^53,54^. CYP treated mice display voiding disfunction, vesical pain, and hematuria that mimics IC symptoms in human patients ^53,54^. The etiology of IC is often unknown and historically it has been defined by the presence of sterile urine. However, recognizing that standard clinical microbiological lab culture conditions cannot detect many common urogenital microbes, several studies have searched for a potential microbial cause of IC ^55-57^. A few reports have linked *Gardnerella* to IC. The earliest was a small study in 1989 that detected *Gardnerella* in catheterized urine and bladder biopsies from patients with IC ^58^. Recently one study found increased *Gardnerella* in Hunner lesion IC compared to non-Hunner IC ^59^. However, another study did not detect *Gardnerella* by PCR in bladder biopsies from patients with IC, albeit using formalin-fixed parafin-embedded tissue ^60^. It is important to note that all these studies sought to detect bacteria in urine or bladder tissue at the same time of symptoms and pathology. However, most strains of *Gardnerella* that caused edema and exfoliation in mice behaved as “covert pathogens” that did not persist in the urinary were not detectable at the time the pathology was observed. Our observation that *Gardnerella* strains largely do not persist in the urinary tract suggests that associations may be missed in clinical studies. *Gardnerella* could be initiating mucosal damage but be absent by the time symptoms develop. In future studies, examining whether *Gardnerella* is colonizing adjacent niches like the vagina or periurethral space that could seed transient urinary tract exposures is crucial not only for defining its association with diseases like IC, UUI, and UTI but also for targeting future therapies to the appropriate niche.

The focused nature of this first examination of *Gardnerella* uropathogensis leaves many open questions and opportunities for future studies. Here bladder pathology was examined at a single timepoint following two inoculations. Therefore, it is not yet known how long the urothelial disruption and inflammation persists. Bladder infection by UPEC triggers exfoliation and inflammation that results in epithelial remodeling and epigenetic changes that persist long after the UTI resolves ^61,62^. Whether similar remodeling happens or the urothelium returns to homeostasis following *Gardnerella* exposures remains to be determined. It is possible that more frequent *Gardnerella* exposures would reveal uropathogenic features of additional strains or enhance the ability of a strain to persist in the urinary tract. Clade 3 strains were not equally represented among the strains tested *in vivo*, and therefore the study could be underpowered to detect clade 3 pathogenicity. However, clade 3 strains are less common in the literature ^11^, and our genomic analysis shows that this clade lacks most of the genes that were associated with urinary tract colonization and pathogenesis. Here each *Gardnerella* strain was tested individually. Multiple *Gardnerella* clades often coexist in the vaginal microbiome. The extent to which this is also true in the urobiome has not been investigated. Women with BV typically have a higher diversity and abundance of *Gardnerella* species compared to those with *Lactobacillus*-dominated microbiomes. Future work could examine whether *Gardnerella* clades have combinatorial effects in the urinary tract. This study used young female wild type C57Bl/6 mice. The literature suggests that host factors such as age, menopause status, and diabetes could affect urobiome composition ^63-65^. Some studies have reported that *Gardnerella* was more frequently cultured or detected in urine from older, post-menopausal women ^21^. Therefore, it is possible that colonization is host context specific. Future studies in aged and ovariectomized mice would provide important insights into the colonization potential and temporal dynamics of *Gardnerella* strains from different clades in different urinary tract environments. Similarly, the degree to which *Gardnerella* virulence patterns are shared or differ between the urinary tract and the vagina remains to be determined.

It has been more than a decade since the proposition that *Gardnerella* is a genus composed of at least four distinct clades and several years since new species designations, including *G. vaginalis, G. piotii, G. picketii, G. swidsinskii, G. leopoldii*, and *G. greenwoodii* were proposed ^5,66^. Correlations between distinct clades and BV status, *in vitro* phenotypes, and metronidazole resistance have been observed. Observable differences in potential virulence gene carriage (e.g. vaginolysin, sialidase) and virulence attributes (biofilm formation) have lent support for the hypothesis that certain *Gardnerella* are more pathogenic than others. In this *in vivo* strain-level comparison of *Gardnerella* pathogenesis, we provide direct evidence of differential pathogenicity in the urinary tract and identify genetic signatures related to distinct *in vivo* phenotypes. The comparative genomics results open the door for future *in vivo* comparisons to tease apart the necessity of the genes identified here for *Gardnerella* persistence and pathogenesis in the urinary tract. The finding of clade and strain-level differences in uropathogenicity among *Gardnerella* sets the stage for futures studies to define the molecular pathways through which *Gardnerella* induces exfoliation and to determine the relationship between *Gardnerella* genetic diversity and urothelial exfoliation, BV, UTI, and other urologic syndromes.

## ETHICS STATEMENT

The animal study was reviewed and approved by the Institutional Animal Care and Use Committee (IACUC) of Washington University School of Medicine.

## AUTHOR CONTRIBUTIONS

LK and SW performed mouse infection experiments. LK performed immunofluorescent staining. LK and NG performed blinded exfoliation and edema scoring. RFP performed all genomic analyses. LK, NG, and RFP analyzed the data and drafted the manuscript. All authors contributed to the article and approved the submitted version.

## FUNDING

This work was supported by funding from the NIH National Institute of Diabetes, Digestive, and Kidney Diseases (NIDDK) R03DK132442 (N.M.G.) and R01DK137964 (N.M.G.).

## Supplementary Material

**Supplementary Figure S1:**
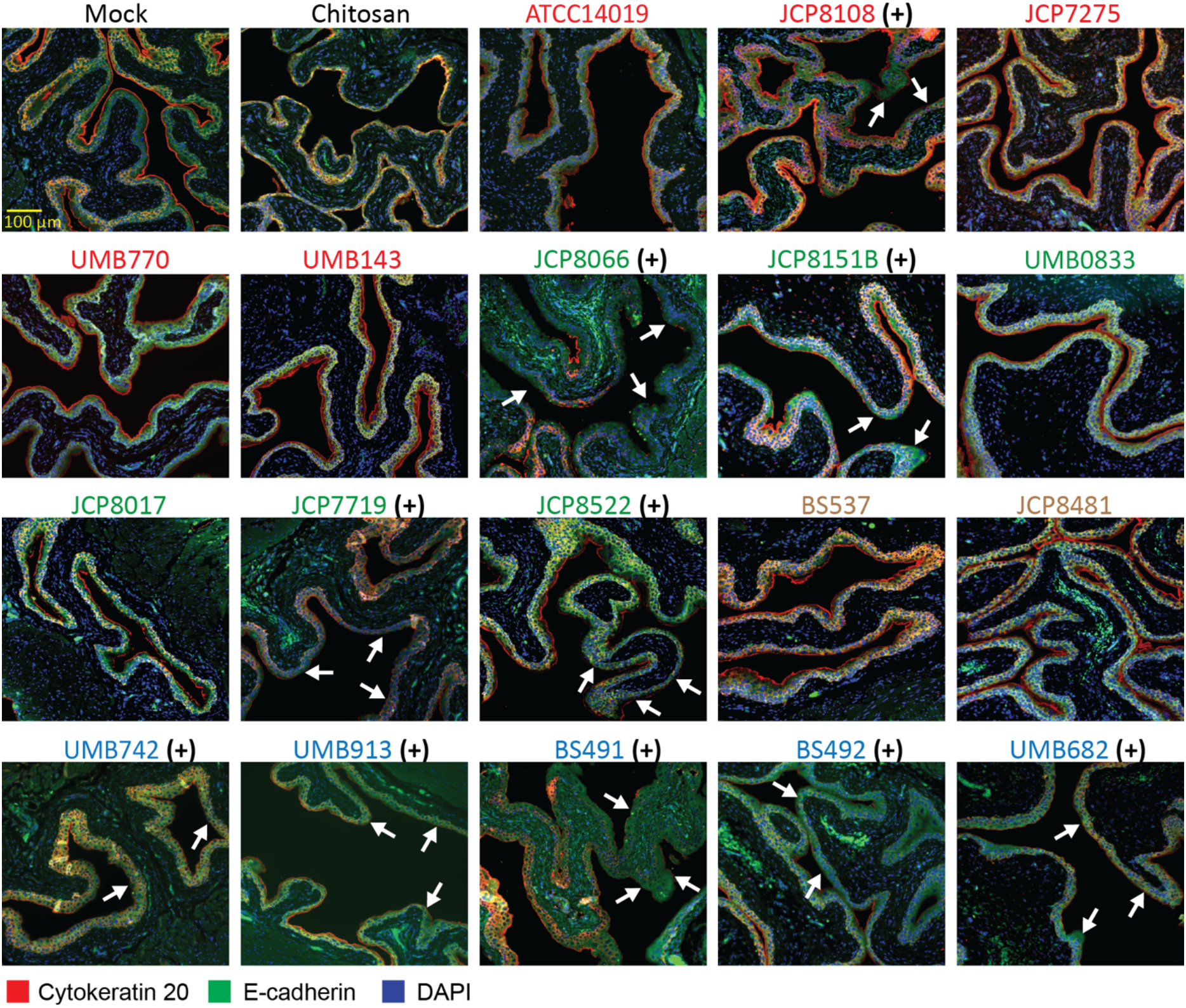
Strain wise exfoliation of urothelium in mouse urinary bladder following *Gardnerella* exposure: Immunostaining images of mouse urinary bladder sections collected after two transurethral exposures of *Gardnerella*. DAPI (blue), e-cadherin (AF488, green) and cytokeratin-20 (AF647, red; marker of terminally differentiated superficial umbrella cells lining the bladder lumen). Areas of exfoliation are denoted by white arrows. Panels display bladder sections from Mock (PBS), Chitosan-treated mice (a transient urothelial disruptor serving as a positive control), and 18 tested *Gardnerella* strains. Scale bar: 100 μm. (+) denotes “Exfoliator” strains based on the statistical analysis shown in **Fig 2B**.

**Supplementary Figure S2:**
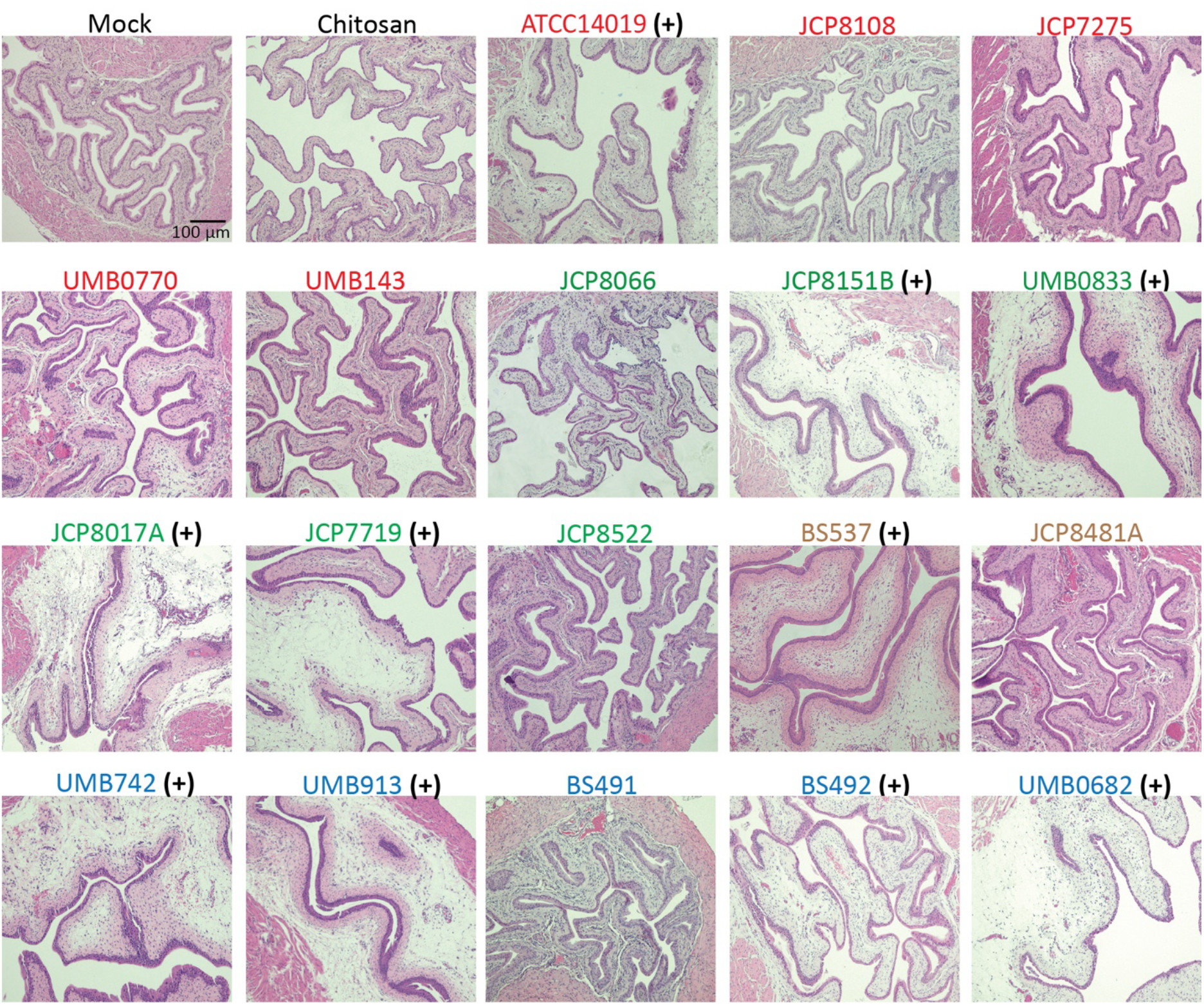
Strain differences in *Gardnerella*-induced edema in the mouse urinary bladder. Representative hematoxylin and eosin (H&E)–stained sections of mouse urinary bladder tissues collected after two transurethral exposures of *Gardnerella*. Panels display bladder sections from Mock (PBS), Chitosan-treated mice (a transient urothelial disruptor), and 18 tested *Gardnerella* strains (Scale bar: 100 μm). (+) denotes “Edema” strains based on the statistical analysis shown in **Fig 3B**.

**Table S1**. Detailed characterization of the 45 associated genes from panaroo, scoary, eggNOG mapper, and BLASTP.

**Table S2**. Detailed information on gene presence/absence from the 291 genome *Gardnerella* pan-genome characterization.

## Notes

### Competing Interest Statement

The authors have declared no competing interest.

